# Redox-mediated inactivation of the transcriptional repressor C3600 makes uropathogenic *Escherichia coli* exquisitely resistant to reactive chlorine species

**DOI:** 10.1101/2021.08.31.458474

**Authors:** Sadia Sultana, Kennadi LeDoux, Mary E. Crompton, Olivia Jankiewicz, Grace H. Morales, Colton Johnson, Elise Horbach, Kevin Pierre Hoffmann, Pooja Kr, Ritika Shah, Greg M. Anderson, Nathan T. Mortimer, Jonathan E. Schmitz, Maria Hadjifrangiskou, Alessandro Foti, Jan-Ulrik Dahl

**Affiliations:** School of Biological Sciences, Illinois State University, Microbiology, Normal, IL, USA; Department of Pathology, Microbiology and Immunology, Division of Molecular Pathogenesis, Vanderbilt University Medical Center, Nashville, TN, USA; School of Biological Sciences, Illinois State University, Cellular Immunology, Normal, IL, USA; Vanderbilt Institute of Infection, Immunology and Inflammation, Nashville, TN, USA; Max-Planck Institute for Infection Biology, Cellular Microbiology, Berlin, Germany

## Abstract

The ability to overcome stressful environments is critical for pathogen survival in the host. One challenge for bacteria is the exposure to reactive chlorine species (RCS), which are generated by innate immune cells as critical part of the oxidative burst. Hypochlorous acid (HOCl) is the most potent antimicrobial RCS and associated with extensive macromolecular damage in the phagocytized pathogen. However, bacteria have evolved defense strategies to alleviate the effects of HOCl-mediated damage. Among these are RCS-sensing transcriptional regulators that control the expression of HOCl-protective genes under non- and HOCl stress. Uropathogenic *Escherichia coli* (UPEC), the major causative agent of urinary tract infections (UTIs), is particularly exposed to infiltrating neutrophils during pathogenesis, however, their responses to and defenses of HOCl are still completely unexplored. Here, we present evidence that UPEC strains tolerate higher levels of HOCl and are better protected from neutrophil-mediated killing compared to other *E. coli*. Transcriptomic analysis of HOCl-stressed UPEC revealed the upregulation of an operon consisting of three genes, one of which encodes the transcriptional regulator C3600. We identified C3600 as a HOCl-sensing transcriptional repressor, which, under non-stress conditions, is bound to the operator and represses the expression of its target genes. During HOCl exposure, however, the repressor forms reversible intermolecular disulfide bonds and dissociates from the DNA resulting in the de-repression of the operon. Deletion of one of the target genes renders UPEC significantly more susceptible to HOCl indicating that the HOCl-mediated induction of the regulon plays a major role for UPEC’s HOCl resistance.

**IMPORTANCE:** How do pathogens deal with antimicrobial oxidants produced by the innate immune system during infection? Uropathogenic *Escherichia coli* (UPEC), the most common etiological agent of urinary tract infections (UTIs), is particularly exposed to infiltrating neutrophils and, therefore, must counter elevated levels of the antimicrobial oxidant HOCl to establish infection. Our study provides fundamentally new insights into a defense mechanism that enables UPEC to fend off the toxic effects of HOCl stress. Intriguingly, the defense system is predominantly found in UPEC and absent in non-invasive enteropathogenic *E. coli*. Our data suggest that expression of the target gene *c3601* is exclusively responsible for UPEC’s increased HOCl tolerance in culture and therefore potentially contributes to UPEC’s survival during phagocytosis. Thus, this novel HOCl stress defense system could potentially serve as an attractive drug target to increase the body’s own capacity to fight UTIs.

## INTRODUCTION

*Escherichia coli* is one of the best and most thoroughly studied free-living organisms. Members of this species are characterized by remarkable diversity: while some *E. coli* strains live as harmless commensals in mammalian intestines, other distinct genotypes represent serious intestinal pathogens that cause significant morbidity and mortality and are categorized into the six distinct pathotypes (1). Yet another group of life-threatening pathogens are extraintestinal *E. coli,* including uropathogenic *E. coli* (UPEC), the most common etiologic agent in approximately 80% of urinary tract infections (UTIs) (2–4). One major difference to intestinal pathogens is that UPEC grow as seemingly harmless commensals in the intestinal environment but rapidly turn into serious pathogens after entry into the urinary tract (5). UPEC ascend from the urethra to the bladder, where they adhere to uroepithelial cells, are internalized and form biofilm-like bacterial communities in the protected intracellular environment of the host cell (2, 6).

However, prior to their attachment to uroepithelial cells, UPEC must surpass host defense mechanisms including phagocytic attack by neutrophils (2). Within the phagosome of neutrophils, bacteria are confronted with a complex mixture of antimicrobial compounds including reactive oxygen and chlorine species (ROS; RCS) (7, 8). Production of neutrophilic ROS and RCS, a process named oxidative burst, involves the assembly and activation of NADPH oxidase. This enzyme complex catalyzes the reduction of molecular oxygen to superoxide in the phagosomal space, which is subsequently dismutated to hydrogen peroxide (H_2_O_2_). Intracellular granules also release myeloperoxidase into the phagosome, an antimicrobial enzyme that converts H_2_O_2_ and available (pseudo-) halides into microbicidal hypohalous acids (9–11). In contrast to H_2_O_2_, which shows only very modest reactivity with most cellular macromolecules and is well tolerated by most bacterial species even at millimolar concentrations (12), hypochlorous acid (HOCl), the most prominent hypohalous acid, is extremely reactive and bactericidal already at low micromolar levels (13, 14). HOCl oxidizes virtually any cellular molecule, including select amino acids, lipids, metal centers, and nucleic acids (15). This, in turn, leads to macromolecular damage and, ultimately, microbial death. One well-known target of HOCl is the amino acid cysteine (8, 9). HOCl or related chloramines oxidize cysteines to either reversible (i.e. sulfenic acids; disulfide bonds) or irreversible oxidative thiol modifications (i.e. sulfinic and sulfonic acid) (16). Reversible thiol modifications often have structural and functional consequences while irreversible thiol modifications lead to protein aggregation and degradation (12, 13).

Bacteria have evolved various strategies to counteract and reduce the harmful effects of ROS/RCS stress. ROS/RCS significantly affect global gene expression, which often is a result of changes in the activities of redox-regulated transcriptional regulators. Posttranslational modifications of redox-sensitive amino acid side chains in these regulatory proteins affect their promoter DNA binding activity and ultimately the expression of the corresponding stress-protective target genes. Although HOCl is one of the most potent industrial and physiological antimicrobials (18), little is known about the cellular consequences of HOCl stress in Gram-negative pathogens. This is particularly surprising given that bacterial responses to the two less reactive bactericidal oxidants H_2_O_2_ and superoxide have been studied in great detail (19–23). Many of their responses involve select transcriptional regulators, which can distinguish between different stressors through oxidation of conserved cysteine residues (24). So far, only three HOCl-sensing transcriptional regulators have been identified in *E. coli*, all of them in the K-12 strain MG1655: HypT, which is activated through methionine oxidation (25); the TetR-family repressor NemR and the AraC-family activator RclR, which represent two transcription factors that use the oxidation status of cysteine residues to sense and respond to HOCl. Oxidation of NemR leads to its dissociation from the promoter, causing de-repression of its target genes (26). In contrast, the transcriptional activator RclR binds its target DNA upon HOCl-mediated cysteine oxidation, leading to a strong and specific activation of the expression of the *rclABC* operon (27).

It is well accepted that neutrophils and the oxidative burst play a crucial role for the clearance of UPEC during UTI (28). Previous studies have identified defense systems against hydrogen peroxide (H_2_O_2_) that positively affect UPEC’s ability to colonize the bladder emphasizing the importance of bacterial oxidative stress defense systems for UPEC pathogenesis (29–31). However, these defense systems are also present in commensal *E. coli*, which limits their suitability as UPEC-specific drug targets. Here, we demonstrate for the first time that resistance to the most abundant neutrophilic oxidant, HOCl, and the oxidizing environment of the neutrophil phagosome is significantly higher in UPEC compared to other *E. coli* pathotypes indicating the presence of additional HOCl defense systems. Intriguingly, our knowledge on UPEC’s HOCl defense systems is quite limited. Our transcriptomic data show that UPEC respond to sublethal HOCl-stress with the upregulation of an operon harboring three uncharacterized genes that are not present in the commensal *E. coli* strain MG1655. We identified one of them as a HOCl-sensing transcriptional repressor that reversibly loses its repressor activity during HOCl-stress. The thiol-based inactivation mechanism of C3600 is based on intermolecular disulfide bond formation, which likely results in conformational changes and disables the repressor binding to the promoter DNA. C3600’s inactivation results in the de-repression of the downstream targets, one of which we identified as a major contributor to the increased HOCl resistance of UPEC strains.

## RESULTS

### UPEC shows increased growth and survival during HOCl stress

In the bladder, UPEC is confronted with an onslaught of host defense mechanisms including exposure to neutrophilic oxidants (32). Given that elevated levels of these antimicrobials such as HOCl pose an increased risk for bacteria yet UPEC appears to thrive in this environment, we speculated that extraintestinal *E. coli* may tolerate higher HOCl levels than intestinal *E. coli* pathotypes. To test this possibility, we compared the growth behavior of different *E. coli* strains during HOCl stress. To reduce the possibility that media components react with and potentially quench HOCl, we performed our phenotypic assays in MOPS-glucose (MOPSg) minimal media. We cultivated the two genetically distinct *E. coli* strains MG1655 and CFT073 and monitored their growth for 16 hours (hrs) in the presence and absence of increasing HOCl concentrations. MG1655 is a fecal isolate and was adapted to one of the most robust *E. coli* lab strains (33). CFT073, on the other hand, is an uropathogen that was isolated from the blood of a patient with acute pyelonephritis (34). Exposure of MG1655 and CFT073 to increasing HOCl concentrations resulted in concentration-dependent extensions of their lag phase (LPE) (Fig. 1A). We found that HOCl-mediated LPE are more pronounced in MG1655 indicating a higher HOCl susceptibility of that strain compared to CFT073. We then used the growth curve-based assay to compare the sensitivity of additional *E. coli* strains to sublethal HOCl stress by quantifying their HOCl-mediated changes in LPE. This method has previously been found to be most reproducible for assessing HOCl sensitivity (35). We confirmed the drastically higher HOCl sensitivity of MG1655 while the HOCl concentrations tested had only minor effects on the LPE of CFT073 (Fig. 1B). This is significant given that MG1655 is considered much more HOCl-resistant than most other commensal laboratory *E. coli* strains including MC4100, which lacks a 97 kb region encompassing the complete RclR regulon (36). The HOCl-sensing transcriptional regulator RclR activates the expression of *rclA, rclB*, and *rclC,* all of which contribute to HOCl resistance (27). Our LPE data confirmed that MC4100 is indeed less HOCl-tolerant than MG1655 (Fig. 1B). No significant difference in HOCl sensitivity was observed between MG1655 and the enteropathogenic *E. coli* strain O127:H6, the first *E. coli* pathovar causing infantile diarrhea (37) (Fig. 1B). To exclude the possibility that the higher HOCl resistance of CFT073 is a strain-specific phenomenon, we analyzed the growth behavior of additional UPEC strains during HOCl stress including another common UPEC model strain, the cystitis isolate UTI89 (38), as well as the dominant fluoroquinolone-resistant clone EC958 (39) and two clinical isolates from cystitis patients, VUTI149 and VUTI247 (40). Notably, all five UPEC strains showed similar LPE responses in the presence of HOCl and were substantially more resistant than K-12 strain MG1655 (Fig. 1C). We conclude from these data that the increased HOCl resistance is potentially characteristic for the UPEC pathotype in general. In contrast, MG1655 and CFT073 displayed similar sensitivity to H_2_O_2_, another neutrophilic oxidant generated during phagocytosis, excluding the possibility that UPEC’s increased resistance is targeted to ROS/RCS in general (Fig. S1B).

**FIG 1.**
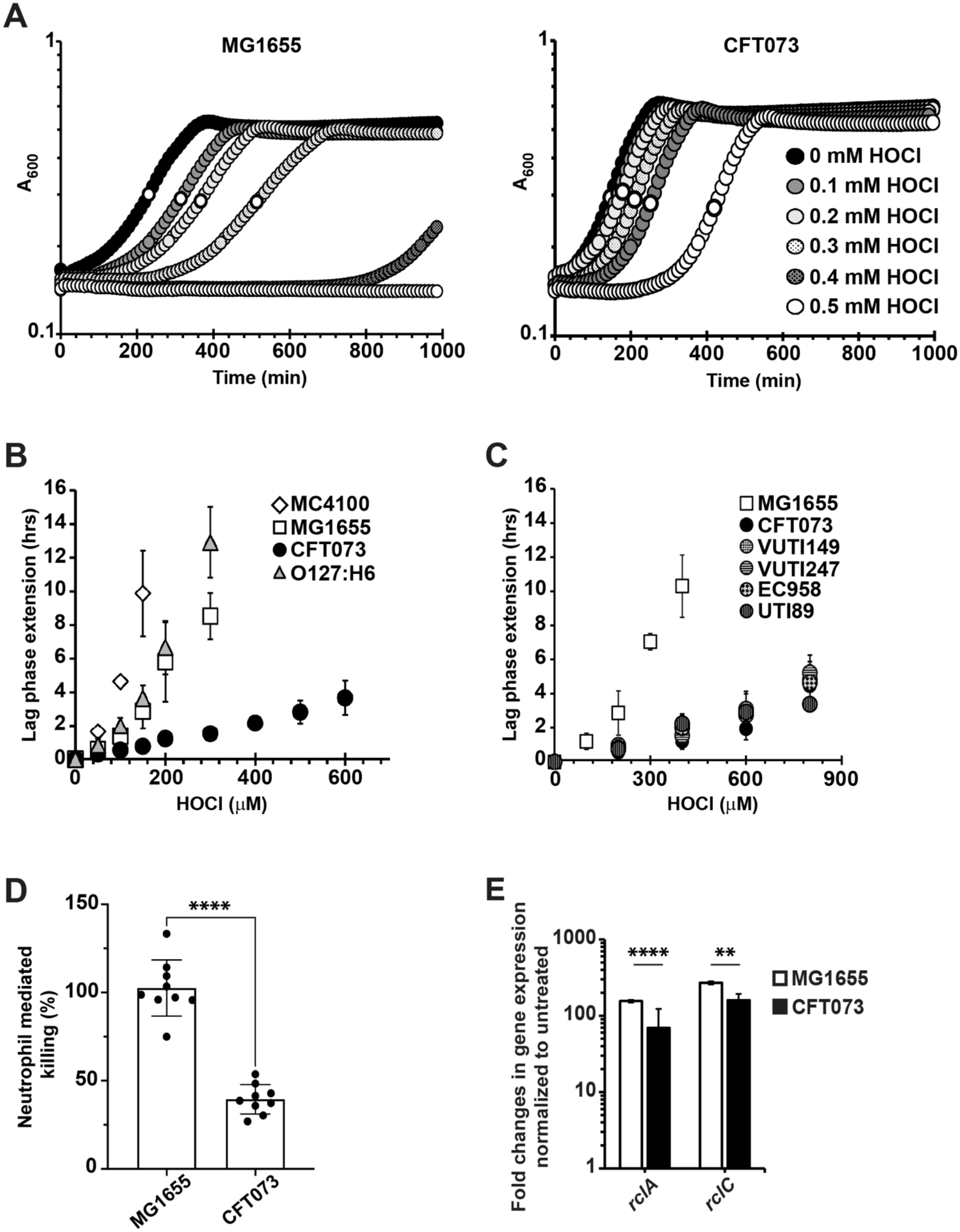
UPEC strains show increased protection from HOCl stress and neutrophil-mediated killing compared to strains from other *E. coli* pathotypes. (A-C) Different *E. coli* strains were cultivated aerobically in MOPSg media in the presence of the indicated concentrations of HOCl. Growth was monitored for 16 hrs at 600 nm. (A) Treatment with sublethal HOCl concentrations causes a concentration-dependent lag phase extension (LPE) in strains MG1655 and CFT073. This effect is less pronounced in CFT073 indicating its increased HOCl resistance. (B; C) HOCl-mediated LPE was calculated for each strain (see Material and Methods for a detailed protocol). (B) LPE of HOCl-stressed *E. coli* strains MG1655 (white squares) and MC4100 (white diamonds) as well as enteropathogenic *E. coli* O127:H6 (grey triangle) were significantly increased compared to UPEC strain CFT073 (black circles). (n=7, ± S.D.). (C) All five UPEC strains (circles) showed higher HOCl resistance than commensal strain MG1655 (white square) (n=3, ± S.D.). (D) Serum-opsonized *E. coli* strains MG1655 and CFT073 were incubated with neutrophils isolated from human blood at MOI of 10:1 for 60 min at 37 °C. Killing of each strain was determined by plating on LB agar for colony forming units. UPEC strain CFT073 was 62% more resistant to neutrophil-mediated killing than commensal *E. coli* strain MG1655, **** *p*<0.0001; (n=3 [with three technical replicates each], ± S.D.). (E) *rclA* and *rclC* expression was determined by qRT-PCR in HOCl-treated CFT073 (black bar) and MG1655 (white bar). The expression of both genes was significantly reduced in CFT073 compared to MG1655. ** 0.01>*p*>0.001; **** *p*<0.0001; (n=3, ± S.D.).

Next, we investigated whether MG1655 and CFT073 also differ in their sensitivity to phagosomal killing by neutrophils, which generate HOCl as the major reactive species during oxidative burst (7). Freshly isolated neutrophils were incubated for 45 min with a 10-fold excess of opsonized MG1655 and CFT073, respectively. Neutrophils and phagocytized bacteria were then separated from non-ingested bacteria and plated for colony forming unit counts after lysis of the neutrophils. Intriguingly, we found that killing of phagocytized CFT073 is ∼62% reduced compared to commensal *E. coli* MG1655 (Fig. 1D) suggesting that a higher HOCl resistance could play a role for UPEC’s ability to survive neutrophil infiltration and establish disease.

### HOCl causes extensive transcriptional changes in UPEC

In contrast to *E. coli* MG1655, the HOCl stress response of UPEC strains is still completely unexplored. As a first test to determine whether HOCl-stressed CFT073 and MG1655 share similarities in gene expression, we determined the sublethal HOCl concentrations for each strain and then used quantitative real-time PCR (qRT-PCR) to compare the HOCl-induced expression of the *rclC* and *rclA* genes. Both genes are members of the RclR regulon, the expression of which is induced in HOCl-stressed MG1655 (27). We confirmed the transcriptional upregulation of both genes in MG1655 and found that their expression was also significantly induced in CFT073 although the fold-changes in *rclA/rclC* expression were significantly lower compared to MG1655 (Fig. 1E). These data suggest that UPEC may employ additional HOCl defense systems, which are not present in EPEC or commensal *E. coli* and which at least in parts compensate for UPEC’s upregulation of the RclR regulon under HOCl stress.

Next, we conducted RNAseq analysis to globally monitor changes in CFT073 gene expression in response to sublethal HOCl stress. CFT073’s chromosome shows a mosaic structure in the distribution of backbone genes, which are conserved between MG1655 and CFT073, and “foreign” genes that presumably have been acquired horizontally (5). Genomic comparisons of different UPEC strains including CFT073 revealed that they share more than 80% of their open reading frames but only approximately 40% of their genome is also found in commensal K-12 strain MG1655 (34). We therefore reasoned that any CFT073 genes that are highly up-regulated upon HOCl stress and that are either not present or not significantly overexpressed in HOCl-treated MG1655 (26) could contribute to CFT073’s elevated HOCl resistance. For the transcriptome analysis, we included all genes annotated in the NCBI database and compared the expression values of the stress treated cells to nonstress treated controls. We set a false discovery rate (FDR) of <0.005 as a threshold for significance and considered transcripts as upregulated when they showed a log2 fold change of >1.5 and downregulated when they showed a log2 fold change of <-1.5. Treatment of CFT073 with 2.25 mM HOCl for 15 min caused the upregulation of 757 genes and downregulation of 681 genes, most of which have an unknown function (Fig. 2A; Table S1). While 32% of these genes are also differentially expressed in HOCl-stressed MG1655 (26), the remaining 68% are either UPEC-specific genes or not induced by HOCl in MG1655 and of particular interest for us. Likely due to the proteotoxic nature of HOCl (41), members of the heat shock response were among the most upregulated genes in HOCl-stressed CFT073 cells, including genes that encode proteases and the molecular chaperones IbpA, IbpB, Spy, and HslO/Hsp33, respectively (Fig. 2B). Not surprisingly and consistent with previous studies we also identified antioxidant (e.g. *grxA*; *trxC; ahpCF*) and copper resistance systems (e.g. *cusABCX; cueO*) in the group of highly upregulated genes (19, 26, 35). Moreover, the expression of four previously characterized HOCl-stress defense systems was increased in CFT073 upon HOCl exposure (i.e. *rclRABC, nemRA, cnoX, yedYZ*) (26, 27, 42–44). Our RNAseq analysis also revealed the upregulation of many biofilm genes in HOCl-stressed CFT073 (Fig. 2B; Table S1). These include the curli-producing *csgABC* and *csgEFG* operons, the diguanylate cyclase encoding gene *ydeH*, the biofilm stress resistance gene *ycfR,* and the *pgaABCD* operon, which produces the major adhesin poly-β-1,6-N-acetyl-glucosamine (poly-GlcNAc) and represents a significant contributor to UPEC’s virulence *in vivo* (45, 46).

**FIG 2.**
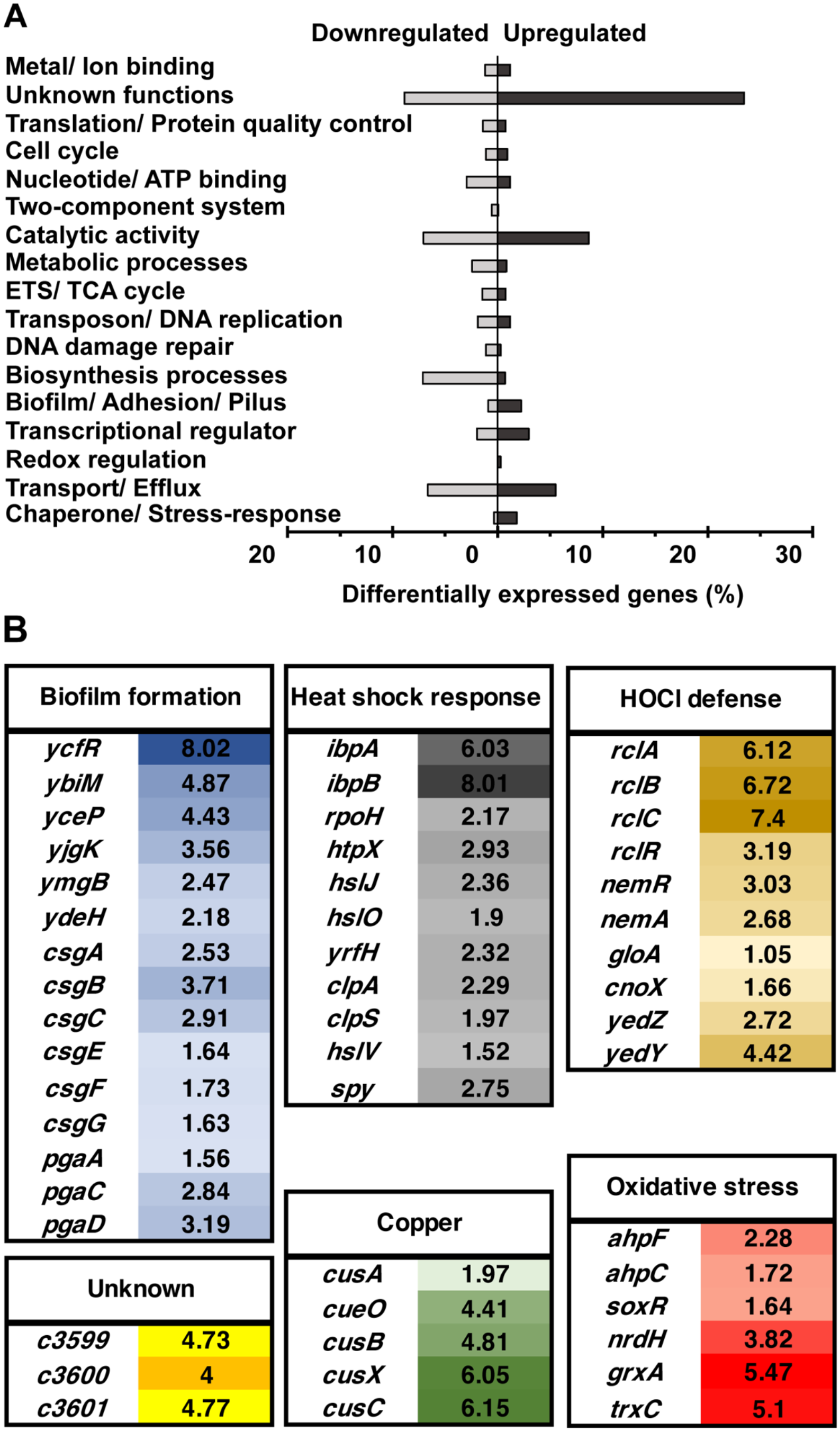
Global gene expression changes in UPEC strain CFT073 in response to sublethal HOCl stress. Exponentially growing CFT073 cells were incubated with a sublethal concentration of HOCl (2.25 mM) for 15 min. Transcription was stopped by the addition of ice-cold methanol. Reads were aligned to the CFT073 reference genome (Accession number: AE014075). (A) Proportion of upregulated (black bars) and downregulated (grey bars) genes of HOCl-stressed CFT073 cells are grouped based on their biological function. (B) Log2-fold change of select CFT073 gene expression at 15 min after sublethal HOCl treatment relative to untreated sample sorted by biological function. The color intensities correlate with the degree of upregulation in gene expression.

### The CFT073 C3600 regulon confers resistance to HOCl stress

32.4% of the differentially expressed genes identified in our RNAseq analysis are uncharacterized and their biological function is still unknown, including a gene cluster consisting of the three genes *c3599*, *c3600*, and *c3601* (Fig. 2, Table S1). The *c3599-c3600-c3601* gene cluster is divergently transcribed from the genes located directly upstream and downstream (i.e. *c3597* and *c3602,* respectively) (Fig. 3A). *c3600* is located downstream of *c3599* and upstream of *c3601* and encodes a putative TetR-family transcriptional regulator, however, the functions of the *c3599* and *c3601* genes are unknown (Fig. 3A). qRT-PCR analysis of HOCl-stressed CFT073 cells confirmed that the transcript levels of these genes are indeed elevated (Fig. S2A). To determine whether C3600 also functions as a transcriptional repressor and to identify potential genes under its control, we performed gene expression studies in CFT073 wild-type and *Δc3600* cells that were grown to mid-log phase under non-stress conditions. qRT-PCR analysis revealed that in comparison to CFT073 wild-type, expression of *c3599* was 105-fold increased while *c3601* mRNA level were 24-fold higher in the absence of the *c3600* gene (i.e. Δ*c3600*) (Fig. 3B). These data indicate that C3600 represses both genes under non-stress conditions and that the HOCl-mediated upregulation of *c3599/c3601* likely depends on C3600’s dissociation from the promoter resulting in the transcription of both genes. In contrast, expression of *c3597* and *c3602* was not affected by the presence or absence of *c3600* (Fig. 3B). RNA-seq analysis of Δ*c3600* cells under non-stress conditions confirmed the constitutive expression of *c3599* and *c3601* and suggest that C3600 also regulates the expression of several genes involved in sulfur metabolism (Table S2).

**FIG 3.**
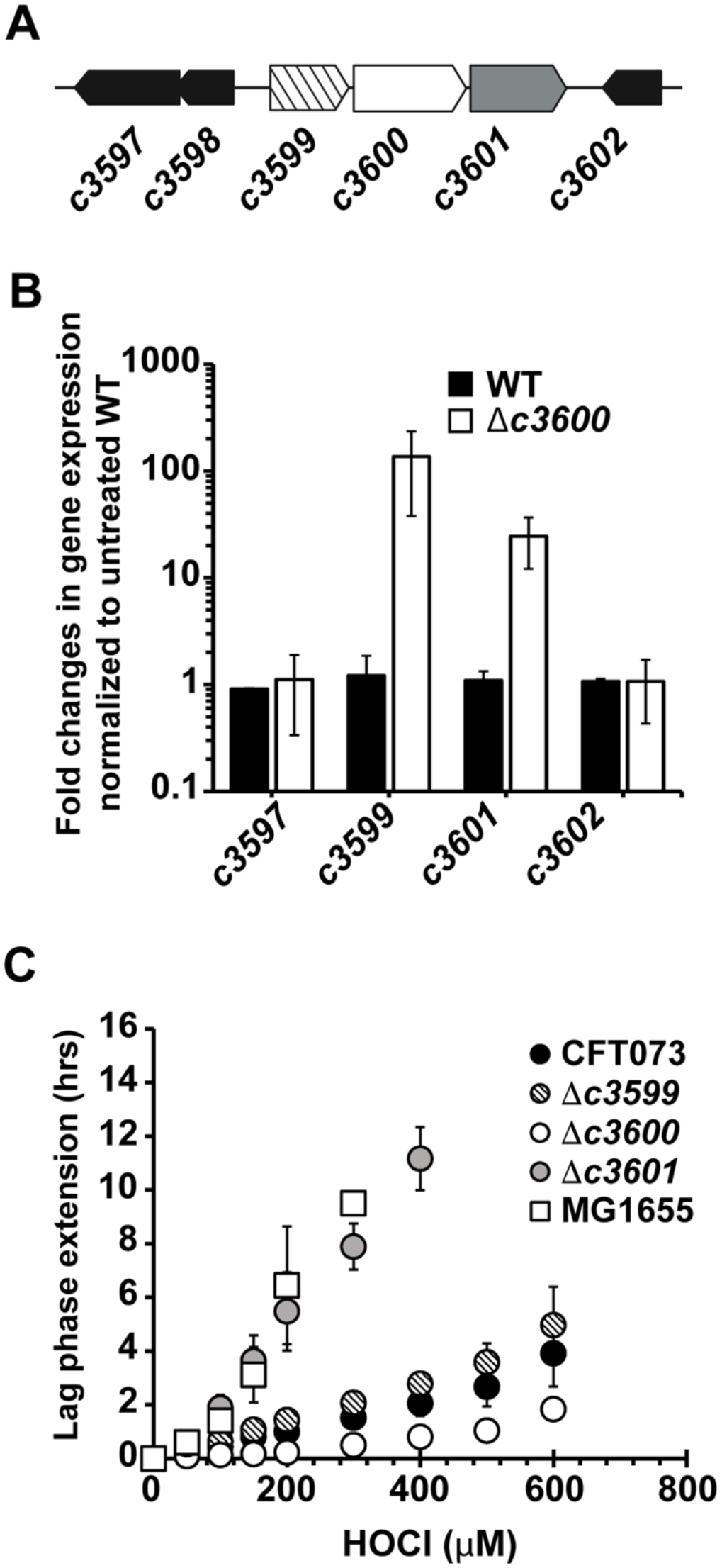
The UPEC-specific gene cluster consisting of *c3599*, *c3600*, and *c3601* protects CFT073 from HOCl stress. (A) Illustration of the gene region containing the *c3599-c3600-c3601* loci. (B) CFT073 wildtype (black bars) and Δ*c3600* cells (white bars) were grown to mid-log phase under non-stress conditions and changes in the expression of the indicated genes were determined by qRT-PCR. *c3599* and *c3601* mRNA level were elevated in the Δ*c3600* strain while expression of *c3597* and *c3602* remained unaffected (n=3, ± S.D.). (C) Growth phenotype analyses of UPEC strains CFT073 wildtype, Δ*c3599*, Δ*c3600*, Δ*c3601,* and commensal *E. coli* strain MG1655 were performed in MOPSg media in the presence of the indicated HOCl concentrations. HOCl-mediated LPE was calculated for each strain (see Material and Methods for a detailed protocol). Deletion of the *c3600* gene rendered CFT073 more resistant whereas *c3601*-deficient cells were substantially more sensitive to HOCl compared to the wildtype (n=5, ± S.D.).

Next, we determined whether any of the three genes play a role for CFT073’s HOCl resistance. We constructed strains with individual in-frame gene deletions and compared their HOCl sensitivities to wild-type cells using the LPE assay. Deletion of *c3599* (i.e. Δ*c3599*) did not significantly affect the LPE (Fig. 3C). However, we found that CFT073 cells that lack the *c3601* gene (i.e. Δ*c3601*) were highly susceptible to HOCl (Fig. 3C). Remarkably, the Δ*c3601* strain was equally sensitive to HOCl as the commensal *E. coli* strain MG1655 suggesting that expression of *c3601* is the main contributor to UPEC’s increased HOCl tolerance *in vitro*. In contrast, deletion of the *c3600* gene (i.e. Δ*c3600*) caused a significant decrease in HOCl-induced LPE indicating that constitutive overexpression of the operon provides additional protection against HOCl (Fig. 3C). Consistently, we found that survival of Δ*c3600* was ∼1 log higher than wild-type CFT073 after 30 minutes exposure to 3 mM HOCl (Fig. S2B). No difference in LPE was observed when CFT073 wild-type and *Δc3600* were exposed to H_2_O_2_ (Fig. S2C).

We searched for the presence of the operon in the genomes of 196 *E. coli* strains from eight distinct pathotypes. While the *c3599-c3600-c3601* gene cluster has not been found in enterohemorrhagic *E. coli* (EHEC), enteropathogenic *E. coli* (EPEC), and enterotoxigenic *E. coli* (ETEC) strains, it was present in 25% of adherent-invasive *E. coli* (AIEC), 8% of avian pathogenic *E. coli* (APEC), 10% enteroaggregative *E. coli* (EAEC), 22% of commensals, and 46% of UPEC (Fig. 4; Fig S3, Table S3). Intriguingly, the exquisitely more HOCl-sensitive intestinal *E. coli* strains MC4100, MG1655 and O127:H6 (Fig. 1B) lack the presence of the *c3599-c3600-c3601* gene cluster. Taken together, our results indicate that we have discovered a novel HOCl stress defense system, which is present in multiple *E. coli* pathotypes and consists of the HOCl-sensing transcriptional repressor C3600 and its regulatory target C3601, an important contributor to UPEC’s HOCl resistance.

**FIG 4.**
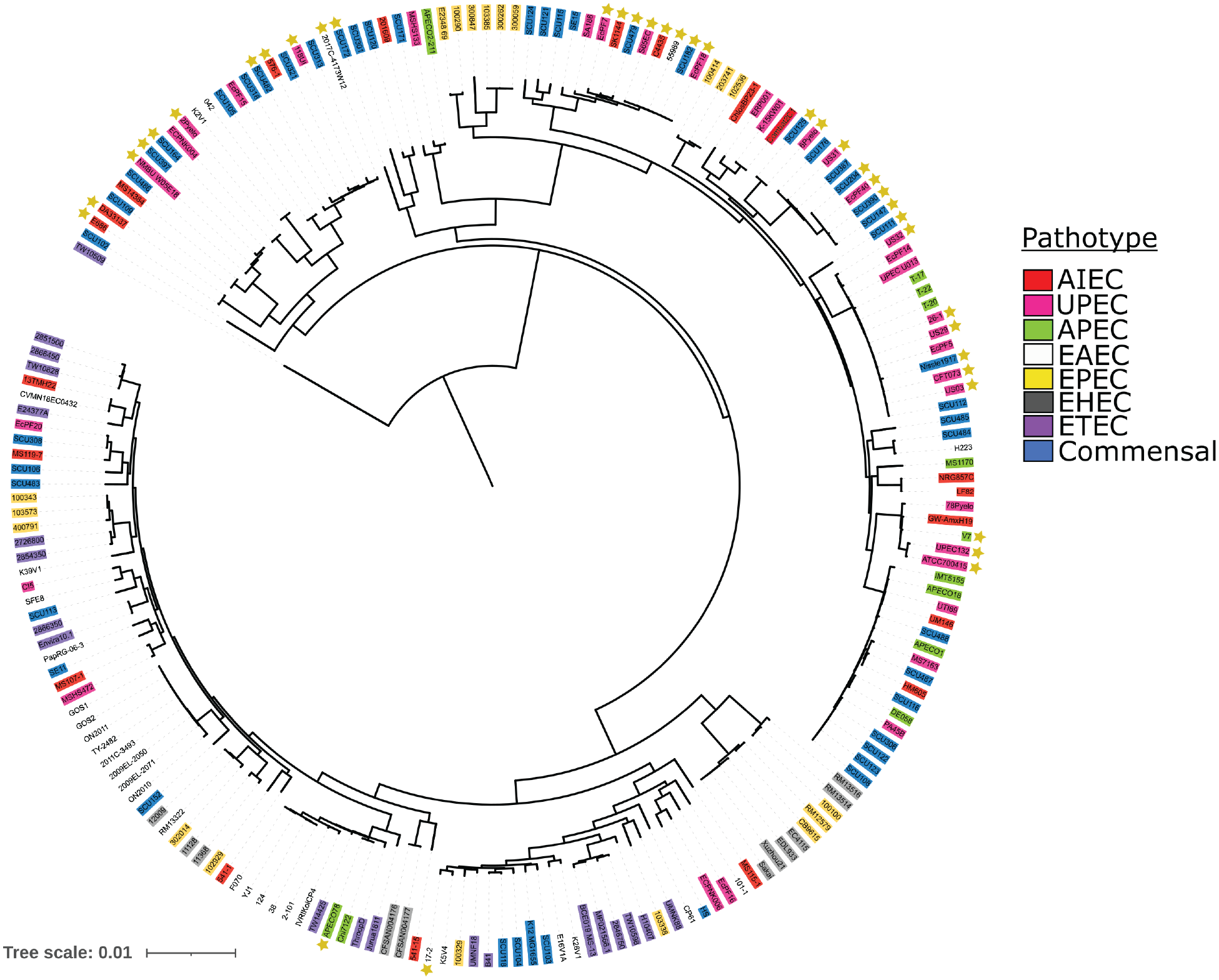
Distribution of *c3599-c3600-c3601* operon in different *E. coli* pathotypes. Genomes from 196 *E. coli* strains of eight pathotypes were downloaded from NCBI and a custom BLAST database was created. A core genome alignment was constructed using Roary version 3.13.0, and a maximum likelihood tree built using IqTree version 2.1.4_beta. The tree was visualized and annotated using Interactive Tree of Life with the pathotype and operon presence. *E. coli* pathotypes used for comparison: AIEC, adherent-invasive *E. coli*; APEC, avian pathogenic *E. coli*; EAEC, enteroaggregative *E. coli;* EHEC, enterohemorrhagic *E. coli;* EPEC, enteropathogenic *E. coli*; ETEC, enterotoxigenic *E. coli*; commensal *E. coli*; UPEC, uropathogenic *E. coli*. The yellow star indicates the presence of the operon, which was lacking in EHEC, EPEC, and ETEC and present in 25% of AIEC, 8% of APEC, 10% EAEC, 22% of commensals, and 46% of UPEC.

### DNA binding of C3600 is inhibited by reversible thiol oxidation under HOCl stress *in vitro*

To assess the ability of C3600 to bind to the operator sequence upstream of the *c3599* gene, we purified wild-type C3600 and conducted *electrophoretic mobility shift assays* (EMSA) using a DNA fragment that contains the promoter region of the *c3599-c3600-c3601* operon. We found that reduced wild-type C3600 binds with high affinity to the *c3599-c3600-c3601* upstream region *in vitro* (Fig. 5). Next, we analyzed the DNA-binding ability of C3600 after oxidation with *N*-chlorotaurine (NCT), a mild, long-lived oxidant generated in innate immune cells as a result of HOCl’s reaction with taurine (13). We used NCT here to limit the risk of protein carbonyl formation and protein aggregation (47). NCT treatment substantially decreased the DNA-binding activity of C3600 (Fig. 5), which could be reversed with the thiol-reducing agent DTT (Fig. S4A) suggesting that C3600 senses RCS stress via a reversible oxidative cysteine modification. Not surprisingly and consistent with our *in vivo* data (Fig. S1C; Fig. S2C), pre-treatment with H_2_O_2_ had no effect on C3600’s DNA-binding ability (Fig. S4B).

**FIG 5.**
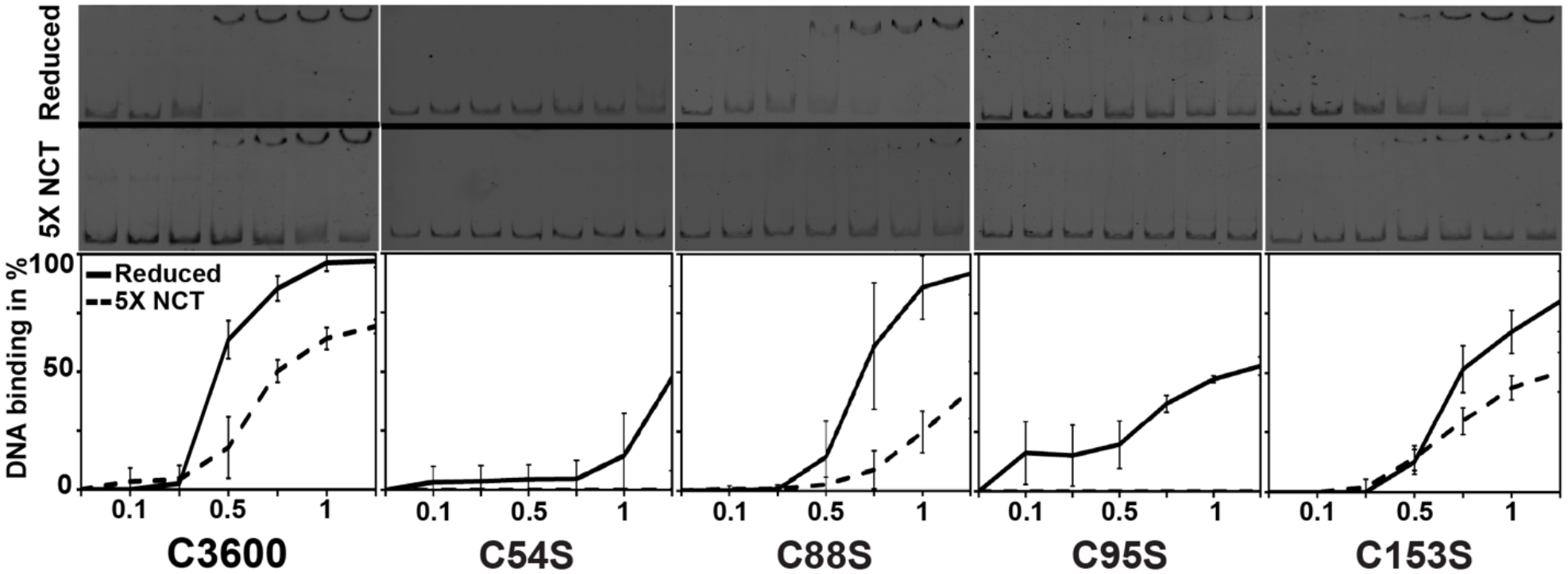
The DNA binding activity of C3600 is inhibited by thiol oxidation after treatment with RCS *in vitro*. Binding of C3600 protein variants (0-1.25 μM) to the *c3599-c3600-c3601* promoter region (P*c3599*, 2 ng) in their reduced states (RED; solid line) and after treatment with a 5-fold excess of N-chlorotaurine (NCT; dashed line) was analyzed in EMSA assays. DNA-binding was visualized by 6% TBE-PAGE and assessed by densitometric quantification using ImageJ. DNA binding ability of all the variants was reduced upon oxidation with NCT. DNA binding of C3600 variants C54S and C95S was impaired even in their reduced state and completely abolished after NCT treatment. Representative gels are shown along with the results of the quantification (n≥3, ± S.D.).

To examine the role of C3600’s four cysteines in DNA-binding and RCS-sensing, we constructed four His_6_-tagged C3600 variants, in which one of four cysteine residues is individually replaced by serine (i.e. C54S, C88S, C95S, C153S, respectively), and tested their DNA binding activity in the presence and absence of NCT. DNA-binding of C3600 variants C88S and C153S were comparable to the wild-type protein (Fig. 5). In contrast, the binding of the variants C54S and C95S was severely compromised even under reducing conditions and completely abolished when the proteins were pre-treated with NCT suggesting that replacement of these cysteines may cause conformational changes that negatively affect DNA-binding. These results support our *in vivo* studies and indicate that RCS-mediated oxidation of C3600 causes its dissociation from the promoter and results in the de-repression of *c3601/c3599*.

### C3600 responds to HOCl stress by intermolecular disulfide bond formation

Given the reversible nature of C3600’s dissociation from the promoter, we excluded the possibility that irreversible cysteine modifications such as sulfinic (-SO_2_H) and sulfonic acids (-SO_3_H) are formed during RCS treatment. We then wondered whether the RCS-mediated inactivation of C3600’s repressor activity is a result of intermolecular disulfide bond formation, a reversible cysteine modification associated with redox-signaling. We treated purified C3600 wildtype and the four variant proteins with 5- and 10-molar excess of HOCl prior to separation by non-reducing SDS-PAGE. In the absence of HOCl, all protein variants migrated primarily in their monomeric form (∼21 kDa) although some disulfide-bonded species were detected in untreated C88S and C95S variants, respectively (Fig. 6A). Upon treatment with HOCl, all variant proteins formed disulfide-linked dimers and intermolecularly disulfide-bonded oligomers, which migrated at ∼43, and 65 kDa, respectively, and could be reversed by the addition of DTT. No such disulfide-bonded species were formed after incubation of wild-type C3600 with H_2_O_2_ (Fig. S5A). Based on these data, we hypothesized that a C3600 variant lacking all four cysteine residues (C3600-4C-S) should not be able to form inter-subunit disulfides. Our attempts to purify the C3600-4C-S variant were not successful potentially due to their crucial role in protein folding. To confirm the formation of the intermolecular disulfides in C3600 *in vivo*, we performed thiol trapping experiments. We individually expressed the plasmid-encoded His_6_-C3600 variants in mid-log BL21(DE3) cells, exposed them to HOCl and separated them by non-reducing SDS-PAGE followed by immunodetection of His_6_-C3600. The ∼30 kDa band appears to be unspecific as it was also detected in control cells that only carry the empty vector (Fig. 6B). Disulfide-linked dimer formation (∼43 kDa) was observed for HOCl-treated cells that express wild-type protein (Fig. 6B) as well as in variants with individual cysteine substitutions (data not shown). In contrast, the 4C-S variant did not respond to HOCl treatment and remained monomeric (∼21 kDa) (Fig. 6B). Coomassie-stained non-reducing SDS gels confirmed that the monomeric nature of the 4C-S variant after HOCl-treatment, while the ∼ 21 kDa band disappeared in HOCl-treated cells expressing the wild-type protein (Fig. S5B). Our results indicate that the HOCl-sensing mechanism of C3600 is cysteine-dependent even though none of the cysteine residues appears to play the primary role for disulfide-bond formation under the conditions tested.

**FIG 6.**
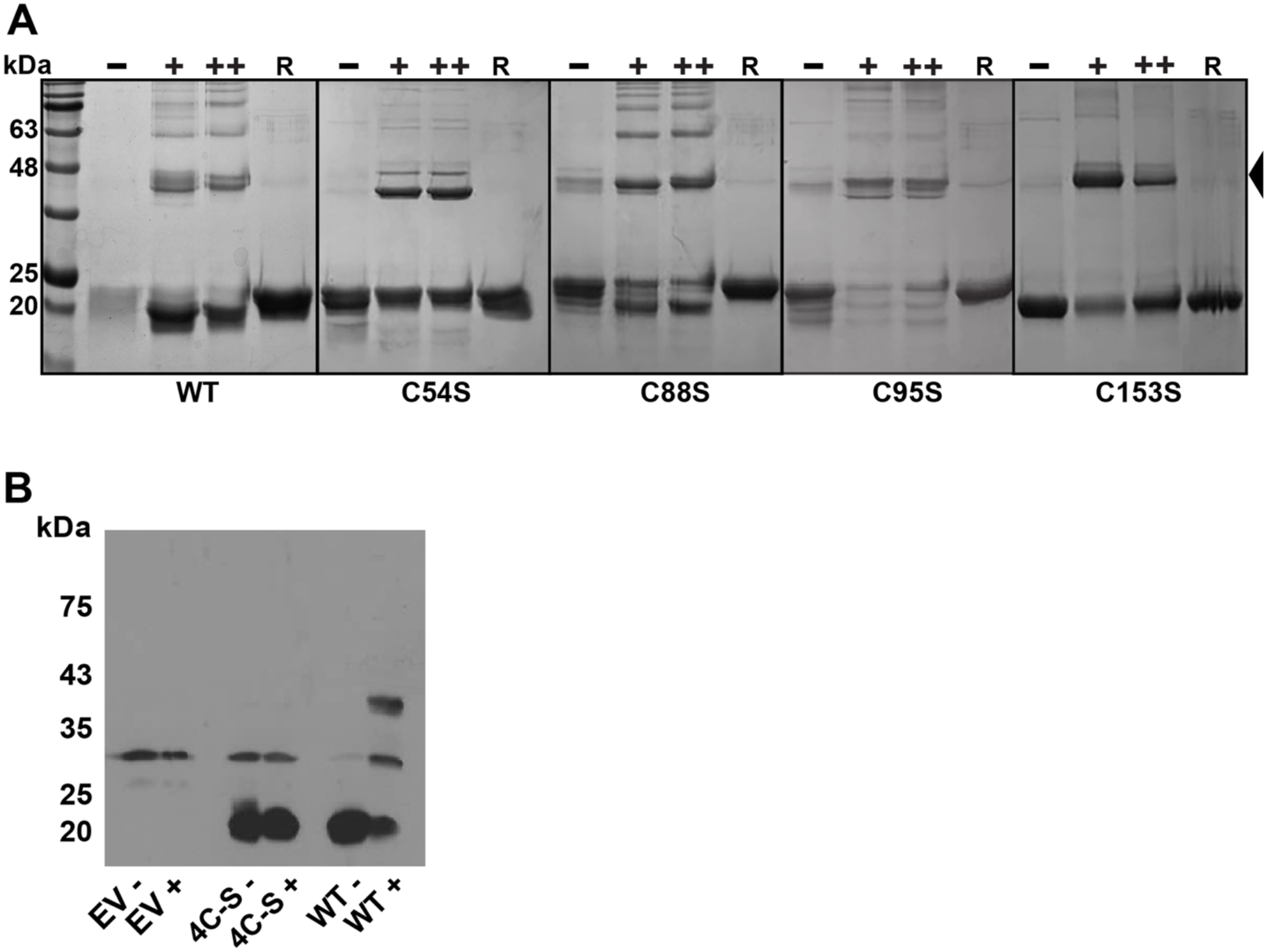
C3600 forms reversible intermolecular disulfide bonds upon exposure to HOCl. (A) 10 μM purified C3600 variants were either left untreated (-) or treated with a 5-(+) and 10-(++) molar ratio of HOCl for 15 min. Proteins were separated by nonreducing SDS-PAGE and visualized by Coomassie staining. The formation of dimers (indicated by the arrow) and higher oligomers was observed in all five HOCl-treated C3600 protein variants. Disulfide bond formation could be reversed by addition of 2 mM DTT (R). Results were verified in four independent experiments. (B) *E. coli* BL21(DE3) expressing His_6_-tagged C3600 variants WT, 4C-S and the empty expression vector (EV) pET28a were grown to mid-log phase, induced with 100 μM isopropyl 1-thio-β-D-galactopyranoside for 60 min, and then either left untreated (-) or treated with 2.5 mM HOCl (+) for another 15 min. Cells were harvested and reduced cysteines irreversibly alkylated with iodoacetamide. C3600 was visualized by Western blotting using nonreducing SDS-PAGE. Results were verified in three independent experiments.

### Cysteine residues 54 and 95 are crucial for DNA-binding and HOCl-sensing *in vivo*

Our *in vitro* DNA-binding assay suggested a potential role for C54 and C95 in redox-sensing. To validate their involvement in C3600’s HOCl response *in vivo*, we performed the LPE-based growth assay using Δ*c3600* cells expressing plasmid-encoded cysteine variants in presence of various HOCl concentrations. Empty vector (EV) and wild-type C3600 complemented strains served as controls. The LPE data fully support our previous findings and showed that expression of *c3600* with serine substitutions in Cys54 or Cys95 were unable to complement the protein function resulting in increased HOCl resistance of these strains like the empty vector control (Fig. 7A). In contrast, expression of the C88S and C153 variants successfully complemented the Δ*c3600* strain.

**FIG 7.**
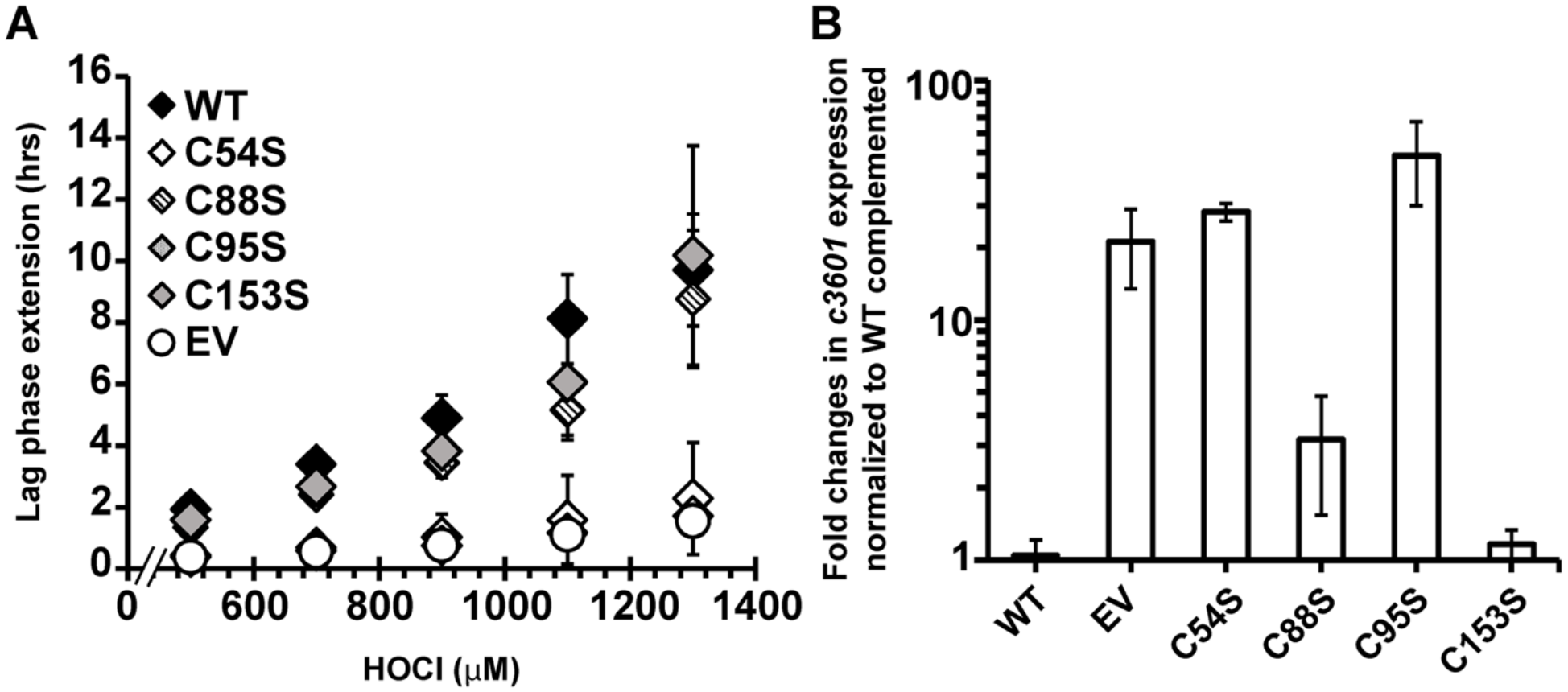
Substitution of Cys54 and Cys95 with serine results in the loss of C3600’s repressor activity in culture. (A) Δ*c3600* strains complemented with plasmids expressing the C3600 variants WT, C54S, C88S, C95S, and C153S, respectively, as well as with the empty vector (EV) pET28a were cultivated in the presence of increasing HOCl concentrations and their LPE was calculated as described in the Material and Methods. (B) The same strains were grown to mid-log phase under non-stress conditions and *c3601* expression was determined by qRT-PCR (n=4, ± S.D.). Expression of C3600-C88S and C3600-C153S complemented the Δ*c3600* strain whereas no complementation was observed for strains expressing C3600-C54S and C3600-C95S (n=4, ± S.D.).

Our transcriptomic analyses revealed that C3600 represses the regulatory target genes under non-stress conditions (Fig. 3B). This phenotype could be reversed by expressing the plasmid-encoded wild-type *c360*0 in Δ*c3600* cells (Fig. 7B). Similarly, expression of the C88S and C153S variants resulted in at least partial complementation of the phenotype. Complementation of Δ*c3600* with plasmids expressing the variant proteins C54S and C95S resulted in an increased expression of *c3601* similar to the EV control, suggesting that both cysteines are also important for C3600’s DNA-binding ability in culture. In summary, our findings indicate that C3600 forms intermolecular disulfide bonds in response to RCS which involves C54 and C95. The crosslinked dimers are unable to bind to DNA and no longer repress transcription of the *c3599-c3600-c3601* operon.

## DISCUSSION

### RCS exposure poses a major threat to bacteria which UPEC counter by the expression of the C3600 regulon

How pathogens regulate their behavior in response to interactions with innate immune cells is fundamental for our understanding of host colonization and the role that bacterial pathogenicity plays in infectious diseases. The production of oxidative stress by the host in response to UPEC has been reported in several independent studies along with the identification of ROS defense systems and their contribution to UPEC pathogenicity (29–31, 48, 49). UPEC face a surge in infiltrating neutrophils after arrival in the bladder (2, 4) and likely experience the highest amount of RCS in the phagosome of activated neutrophils. Particularly in inflammatory environments, infiltrating neutrophils produce the HOCl-generating enzyme MPO at concentrations in the low millimolar range, of which up to 30% leak into the extracellular surrounding (50, 51). Intriguingly, UPEC utilizes the expression of a methionine-rich peptide along with an additional methionine sulfoxide reductase system to scavenge HOCl in the periplasm (52). This unique gene cluster is highly conserved in UPEC pointing to the physiological relevance of RCS-mediated oxidative stress during UTI and the importance for UPEC to efficiently prevent RCS-mediated methionine oxidation (52). It is also possible that dual oxidases (Duox) elevate ROS/RCS level in the bladder, which are members of the NOX family and expressed in several tissues and cell types, including mucosal barrier epithelia and uroepithelial cells of the bladder (53). The enzymes possess a peroxidase homology domain with MPO activity vital for the host immune defense (54) and *Duox* knockdown studies revealed increased bacterial colonization and significantly higher death rates of the hosts (55–57). In fact, RclA, a member of the HOCl-specific RclR regulon, has been shown to protect *E. coli* from Duox-mediated oxidative stress *in vivo* (35). Thus, HOCl production and its ability to control the bacterial population in the host is physiologically relevant (58, 59), potentially providing a rationale why inflammation-associated pathogens such as UPEC have developed strategies to reduce the proteotoxic effects of HOCl.

Treatment with sublethal concentrations of HOCl resulted in a concentration-dependent growth arrest in all *E. coli* strains tested (Fig. 1A-C). HOCl is well known for its high microbicidal activity (16, 51, 60), which explains its role as the active ingredient in household bleach, one of the most commonly used disinfectants in medical, industrial, and domestic settings (18). The oxidant rapidly reacts with a wide range of biological molecules including DNA, lipids, and proteins (9, 15). However, its main mode of killing is mediated through widespread protein unfolding and aggregation as a result of oxidative cysteine and methionine modifications (41). Consistent with previous studies in different bacterial species (19, 26, 61–64), our RNAseq analysis of HOCl-stressed CFT073 revealed the elevated expression of various heat shock genes (Fig. 2; Table S1). The heat shock response is activated as a result of the accumulation of misfolded proteins (65) indicating that proteins are also the major HOCl targets in UPEC.

In comparison to EPEC and commensal *E. coli* strains, all UPEC isolates tested in our LPE analyses were substantially more HOCl resistant (Fig. 1A-C) implying that the UPEC pathotype is generally better equipped to deal with the negative consequences of RCS stress. We identified the *c3599-c3600-c3601* gene cluster as an additional protection system during severe HOCl stress that enables UPEC to grow at higher HOCl concentrations (Fig. 3C). Our bioinformatic search revealed that the operon is predominantly found in invasive *E. coli* pathotypes such as APEC, AIEC and, remarkably, in ∼50% of UPEC strains (Fig. 4; Fig. S3). The different pathotypes are characterized by their specific composition of horizontally acquired genetic material and the individual sets of virulence factors impact their ability to cause disease. It has been proposed that advanced resistance to phagocytosis plays an important role for the pathogenicity of APEC isolates (66). Some APEC strains have been reported to cause UTIs in humans (67), likely due to similarities in important virulence genes present in both pathotypes. It is therefore possible that APEC employ additional RCS stress defense systems, such as the C3600 regulon, to thrive in RCS-rich environments and/or survive infiltrating neutrophils. Indeed, we demonstrated that UPEC strain CFT073 is more resistant to neutrophil-mediated killing compared to the non-pathogenic *E. coli* K-12 strain MG1655 (Fig. 1D). Whether this phenotype is a result of an improved RCS stress defense, potentially mediated by the C3600 regulon, will be subject of our future investigations. Recent studies showed increased cysteine oxidation and oxidative stress in phagocytized *E. coli*, emphasizing the role of HOCl in microbial killing (7, 68). The same study provided strong evidence for HOCl as the main component of the oxidant mixture produced in neutrophils.

Expression of the C3600 regulon is controlled by the transcriptional regulator C3600, which is encoded by one of the members of the operon (Fig. 3A-B). The precise biological function of the other two genes, *c3599* and *c3601,* is still unknown. Deletion of *c3601* resulted in a substantial growth delay comparable to that observed in EPEC and commensal *E. coli* strains (Fig. 3C). This might be due to the inability of the strains to cope with the high level of proteotoxic stress requiring *de novo* synthesis of repair proteins. *c3601* encodes a hypothetical inner membrane protein of the uncharacterized DUF417 protein family that is homologous to RclC, a member of the RclR regulon, which is also expressed in HOCl-stressed UPEC cells (Fig. 1D). The RclR regulon consists of *rclA, rclB*, and *rclC* and is induced through a HOCl-sensing mechanism of the transcriptional activator RclR both in lab culture and during phagocytosis (27, 69). RclA was recently identified as a Cu (II) reductase that protects *E. coli* from HOCl- and Cu-stress (35). While the role of RclB and RclC during HOCl stress is still unknown, a contributing role to RclA’s copper-dependent protective function was proposed. Our future studies are now directed to elucidate the role of C3601 for UPEC’s HOCl stress resistance.

### Redox regulation of the transcriptional repressor C3600

Bacteria have evolved numerous mechanisms on both transcriptional and posttranslational level to fend off the toxic effects that come with the exposure to ROS/RCS. These include the conversion of ATP into the chemical chaperone polyphosphate, which was found to protect UPEC strains from HOCl-mediated protein aggregation (49) and the activation of molecular chaperones such as Hsp33, RidA, and CnoX through thiol oxidation or N-chlorination (41, 42, 70–72). Another level of protection is provided through the transcriptional activation of HOCl stress defense genes, which are directly controlled by oxidative (in-)activation of HOCl-sensing transcriptional regulators (15). One of the best characterized ROS/RCS-sensing regulators is OxyR, which is activated by cysteine oxidation and presents an important virulence factor for the pathogenicity of UPEC strain UCB34 (73). OxyR is prone to oxidation during phagocytosis (68) leading to the elevated expression of the OxyR regulon and contributing to *E. coli*’s ability to evolve resistance to HOCl (74). Similarly, HOCl-mediated intramolecular disulfide bond formation enables binding of the transcriptional activator RclR to the promoter to activate the transcription of the RclR regulon (27). Many of the previously identified bacterial defense systems were significantly higher expressed in our RNAseq of HOCl-stressed CFT073 cells, including Hsp33, CnoX and members of the NemR and RclR regulons (Fig. 2B).

We now add a novel member to the growing list of HOCl-sensing transcriptional regulators in UPEC. Like most redox-sensing transcriptional regulators, C3600 represses the transcription of HOCl-protective genes under non-stress conditions (Fig. 3B). C3600 belongs to the TetR family of transcriptional regulators, which are mostly alpha-helical and active as dimers (75) and shares similarity in both the N-terminal DNA-binding domain and the C-terminal sensing domain with NemR, a broadly conserved HOCl-sensing repressor of the same family that is responsible for the expression of the methylglyoxal-detoxifying enzyme GloA and the N-ethylmaleimide reductase NemR (26). C3600’s RCS-mediated inactivation mechanism is based on reversible cysteine oxidation: in the presence of RCS, C3600 forms intermolecular disulfide bonds both *in vitro* and *in vivo* (Fig. 6). A cysteine-free variant of C3600 no longer responds to HOCl with the formation of disulfide-bonded oligomers (Fig. 6B) further emphasizing that the oxidative inactivation is thiol-based. Likely due to conformational changes during oligomerization, C3600 dissociates from the promoter region (Fig. 5) resulting in the transcriptional upregulation of the *c3599-c3600-c3601* operon (Fig. 7). Deletion of *c3600* resulted in elevated *c3601* and *c3599* mRNA level along with increased resistance to HOCl suggesting that constitutive overexpression of the operon provides additional protection against HOCl (Fig. 3B-C). The DNA-binding activities of C3600 variants C54S and C95S were severely reduced under non stress conditions and completely abolished in the presence of RCS (Fig. 5), providing evidence for the crucial role that both cysteines play for C3600’s HOCl sensing mechanism. This is in contrast to NemR, which uses a completely different set of cysteines for redox-sensing and is inactivated through a thiol:sulfenamide switch (26, 76).

Given that the presence of H_2_O_2_ had no effect on the DNA binding activity, C3600 may very well be HOCl-specific much like the previously characterized transcriptional regulators RclR and HypT (25, 27, 77). This can likely be explained with HOCl’s high reactivity, as it has a 100-times shorter lifetime than H_2_O_2_, acts more local, and rapidly reacts with a variety of biomolecules (9, 24). H_2_O_2_, in contrast, is thiol-specific, orders of magnitude less bactericidal and therefore only kills bacteria after long exposure or at higher concentrations. Probably because they can produce this oxidant as an endogenous byproduct in metabolic reactions (78), bacteria have evolved several efficient systems to eliminate H_2_O_2_ including the peroxiredoxins AhpC and AhpF, whose expression was induced in our RNAseq of HOCl-treated CFT073 (Fig. 2B; Table S1). HOCl, on the other hand, is exclusively produced in eukaryotes and may therefore be perceived by bacteria as a signal for close proximity, which they utilize to adapt their responses (79). We now propose the following model (Fig. 8): under non-stress conditions, C3600 is bound to the operator located upstream of the *c3599-c3600-c3601* operon and represses the expression of the three target genes. Under HOCl stress, however, as it occurs in the phagosome of neutrophils, UPEC senses HOCl through cysteine oxidation of C3600, which forms disulfide-bonded oligomers between Cys54 and Cys95. Likely due to conformational changes, the repressor becomes inactive and dissociates from the operator resulting in the transcriptional upregulation of the *c3599-c3600-c3601* operon.

**FIG 8.**
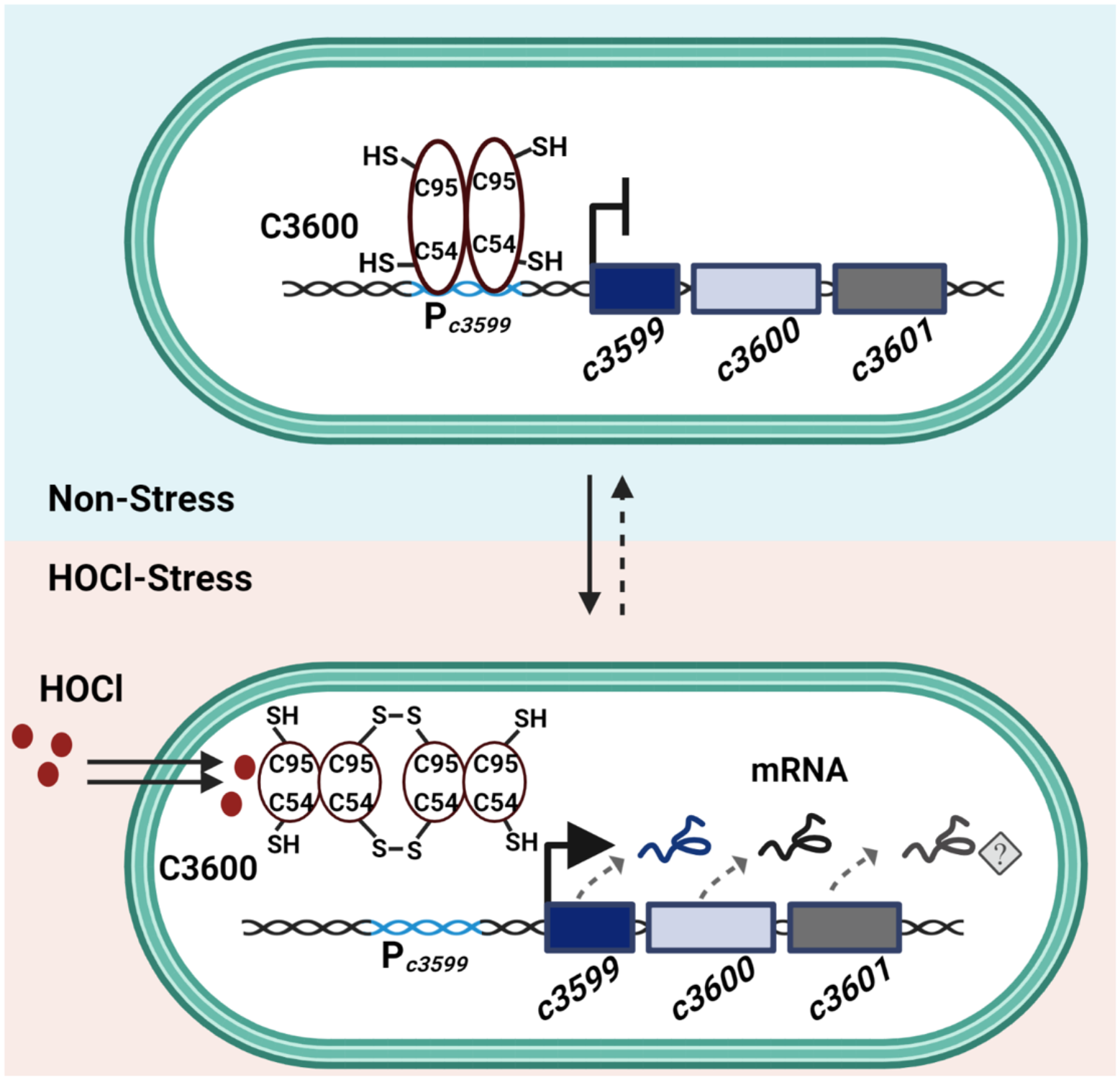
Redox-sensing mechanism of C3600 under HOCl stress. The HOCl-sensing transcriptional repressor C3600 controls the expression of the UPEC-specific genes *c3599, c3600*, and *c3601.* Under non-stress conditions, C3600 is bound to the operator located upstream of the *c3599-c3600-c3601* operon and represses the expression of the three target genes. Under HOCl stress, however, as it occurs in the phagosome of neutrophils, C3600 is oxidized and forms oligomers through inter-subunit disulfide bond formation leading to the inactivation of its repressor function, dissociation from the operator and de-repression of the *c3599-c3600-c3601* transcription. Cys54 and Cys95 are involved in C3600’s redox-sensing and important for the DNA-binding activity of the repressor. Expression of *c3601* contributes to UPEC’s increased resistance to the antimicrobial oxidant HOCl.

### Is a functional HOCl response essential for UPEC colonization and disease?

Many biofilm-forming bacteria respond to changes in the environment, such as exposure to RCS, by switching from a planktonic to sessile growth (80, 81). The lifestyle change provides many survival benefits including up to 1,000-fold increased resistance to antibiotic treatment and protection from clearance even during extensive neutrophil infiltration (82, 83). Our transcriptomic data revealed the HOCl-induced expression of various genes involved in biofilm formation (Fig. 2B; Table S1), including *ydeH*, which encodes a diguanylate cyclase (84). Diguanylate cyclases are responsible for the synthesis of cyclic diguanosine monophosphate (c-di-GMP), a key regulator for biofilm formation that is involved in the regulation of cell surface-associated traits and persistent infections (85). YdeH with its N-terminal Zn^2+^-binding domain has recently been identified as the catalyst for this switch: HOCl-mediated cysteine oxidation disrupts Zn^2+^-binding of the protein and leads to increased diguanylate cyclase activity and elevated c-di-GMP level, which positively affects the production of the exopolysaccharide poly-GlcNAc and facilitates adhesion to bladder cells (79, 86). Poly-GlcNAc production is mediated by the *pgaABCD* operon, which we found to be induced in our RNAseq analysis of HOCl-stressed CFT073 (Fig. 2B; Table S1) suggesting that the YdeH-mediated increase in c-di-GMP indeed activates exopolysaccharide production. Many UPEC isolates express poly-GlcNAc during biofilm formation and host colonization making the polysaccharide a significant contributor to UPEC’s virulence *in vivo* (45, 87, 88). The same Zn^2+^-binding domain and equivalent HOCl-sensing mechanism has been reported in chemoreceptors such as *Helicobacter pylori* TlpD, which facilitates chemoattraction to HOCl sources for *H. pylori* and potentially explains the persistence of this pathogen in inflamed tissue (89). Exposure to HOCl also induced the expression of the curli-producing genes *csgABCEFG* (Fig. 2B; Table S1), which mediate surface attachment and structural integrity in biofilm communities (90) and confer resistance to HOCl treatment (91). To establish disease, UPEC must overcome a plethora of host defenses including neutrophilic attacks before they adhere to and invade uroepithelial cells to form intracellular biofilms (2–4). Moreover, the development of catheter associated UTIs is a result of UPEC’s ability to attach to abiotic surfaces such as catheters in an inflammatory environment. We identified the C3600 operon as a major player for UPEC’s HOCl resistance as it enables growth at otherwise toxic HOCl concentrations (Fig. 3C). The gene cluster is predominantly present in *E. coli* pathotypes that are associated with host cell adhesion and invasion (Fig. 4; Fig. S3). A recent study reported an interdependence of resistance to ROS, biofilm formation and pathogenicity in *Proteus mirabilis*, another leading uropathogen (92). Similarly, EAEC adherence to epithelial cells is stimulated by infiltrating neutrophils presumably due to the presence of additional defense mechanisms to oxidative burst, which are therefore considered beneficial for their pathogenicity (93). However, whether UPEC has evolved the C3600 regulon as a prerequisite for their ability to overcome neutrophilic attacks, thrive in inflammatory environments, and switch from planktonic to sessile lifestyle will be subject of our future investigations.

## MATERIAL AND METHODS

### Strains, plasmids, oligonucleotides, and growth conditions

All strains, plasmids, and oligonucleotides used in this study are listed in Table S4 in the supplemental material. Unless otherwise mentioned, bacteria were cultivated at 37 °C and 300 rpm in luria broth (LB, Millipore Sigma) or in 3-(N-morpholino)propanesulfonic acid minimal media containing 0.2% glucose, 1.32 mM K_2_HPO_4_ and 10 μM thiamine (MOPSg) (94). Kanamycin (100 μg/ml), ampicillin (150 μg/ml) and chloramphenicol (34 μg/ml) were added when required.

### Construction of CFT073 gene deletions

In-frame deletion mutants were constructed using the lamda red-mediated site-specific recombination. CFT073 genes *c3599, c3600*, and *c3601* were replaced with a chloramphenicol resistance (Cm^R^) resistance cassette, which was resolved using pCB20 to yield the nonpolar in-frame deletion strains Δ*c3599,* Δ*c3600*, and Δ*c3601*, respectively (95). All chromosomal mutations were confirmed by PCR.

### Preparation of Oxidants

The molar HOCl concentration was determined by measuring the *A*_292 nm_ of the sodium hypochlorite (Millipore-Sigma) stock solution diluted in 10 mM NaOH using ε_292_ = 350 M^-1^ cm^-1^. The molar H_2_O_2_ concentration was quantified by measuring the *A*_240 nm_ of the stock solution diluted in 50 mM KPi buffer using ε_240_ = 43.6 M^-1^cm^-1^. *N*-Chlorotaurine was prepared by mixing HOCl with excess of taurine (96). All oxidant dilutions were prepared fresh before each use.

### Determining HOCl susceptibility through lag phase extension (LPE) analyses

Overnight LB cultures of the indicated strains were diluted 25-fold into MOPSg and cultivated until late exponential phase was reached. Cultures were diluted into fresh MOPSg to an *A*_600 nm_ = 0.02 and cultivated in a Tecan Infinite 200 plate reader in the presence or absence of the indicated concentrations of HOCl and H_2_O_2_, respectively. *A*_600 nm_ measurements were recorded every 10 min for 16 hours. Oxidant sensitivities of the strains tested were examined by quantifying their oxidant-mediated LPE. LPE were calculated by determining the differences in time for oxidant-treated samples to reach *A*_600 nm_ > 0.3 compared to the untreated controls as described in (35).

### Survival after exposure to HOCl

Cultures of CFT073 and Δ*c3600* were either left untreated or treated with 3 mM HOCl and incubated for 30 min. Excess of HOCl was quenched by adding 5-fold molar ratio of sodium thiosulfate before samples were 10-fold diluted in PBS, spotted onto LB agar plates, and incubated overnight at 37 ⁰C.

### Gene expression analyses by qRT-PCR

Overnight LB cultures of the indicated strains were diluted into MOPSg to an *A*_600 nm_= 0.1 and cultivated until *A*_600 nm_ ∼0.55 was reached before they were either left untreated or treated with 2.5 mM of HOCl for 15 min. 1 ml cells were harvested onto 1 ml of ice-cold methanol to stop transcription. After centrifugation, total RNA was prepared from the cell pellet of three biological replicates of untreated and HOCl-treated CFT073 as well as untreated Δ*c3600*, respectively, using a commercially available RNA extraction kit (Macherey & Nagel). Remaining DNA was removed using the TURBO DNA-free kit (Thermo-Scientific) and cDNA generated using the PrimeScript cDNA synthesis kit (Takara). qRT-PCR reactions were set up according to the manufacturer’s instructions (Alkali Scientific). Expression of the indicated genes was normalized against expression of the 16S rRNA-encoding *rrsD* gene and fold-changes in gene expression were calculated using the ΔΔC_T_ method (97).

### RNA-seq analysis

Samples of HOCl-treated and untreated CFT073 and Δ*c3600* cells were collected as described before for qRT-PCR. After extraction of total RNA (Macherey & Nagel) and removal of the residual DNA using the TURBO DNA-free kit (Thermo-Scientific), rRNA was depleted using the Illumina Ribo Zero Kit (Illumina) for Gram-negative bacteria. 150 base pair single end sequencing was performed on an Illumina HiSeq 2500 by Novogene (Sacramento, USA). Differential gene expression analysis of three biological replicates, including normalization was performed in the bioinformatics platform Galaxy (98). Briefly, RNAseq reads were mapped to *E. coli* CFT073 reference sequence (GCA_000007445.1) using HISAT2 (99). Then the number of reads mapped to each gene were counted using featureCounts (100). Finally, differential gene expression was determined using DESeq2 (101) with an adjusted p-value cut off p ≤ 0.05 and logFC cut off of 1.5.

### Neutrophil-mediated killing of *E. coli*

Human neutrophils were purified from fresh peripheral blood by Histopaque®-1119 (Millipore Sigma) and subsequent discontinuous Percoll® gradient centrifugation (102). Isolated neutrophils were resuspended in RPMI 1640 (without phenol red; Gibco) supplemented with 10 mM HEPES and 0.1% human serum albumin. The rate of bactericidal activity of neutrophils was determined and calculated by a one-step bactericidal assay as described previously (103). 1 x 10^6^ neutrophils were incubated with 1 x 10^7^ opsonized MG1655 or CFT073 in 1 ml of RPMI with 0.1% human serum albumin, with final ratio of bacteria to neutrophils of 10:1. Samples were continuously rotated and incubated at 37 °C for 45 min. Neutrophils and phagocytized bacteria were pelleted by centrifugation at 100 x *g* for 10 min. Supernatants were plated on LB agar plates to determine the number of non-phagocytized bacteria. The pellets containing neutrophils and phagocytized bacteria were washed twice with PBS, lysed in water at pH 11.0, and pelleted by centrifugation at 300 x *g* for 10 min to remove neutrophil debris. The supernatant containing the bacteria was plated on LB agar plates, incubated overnight at 37 °C for CFU counts.

### Phylogenetic tree

Genomes from 196 *E. coli* strains of eight pathotypes were downloaded from NCBI. A custom BLAST database was created locally with these strains. UPEC strain CFT073 genes *c3599*, *c3600*, and *c3601* were identified within the custom database. A core genome alignment was constructed using Roary version 3.13.0 (104), and a maximum likelihood tree built using IqTree version 2.1.4_beta (105). The tree was visualized and annotated using Interactive Tree of Life with the pathotype and operon presence (106). Other graphs were produced using Graphpad Prism 8.4.2.

### Plasmid construction

The *c3600* gene was amplified from UPEC strain CFT073 genomic DNA with primers listed in Table S4 and cloned into the *Nde*I and *BamH*I sites of plasmid pET28a to generate the N-terminally His_6_-tagged C3600 expression plasmid pJUD13. C3600 variant proteins were created using the phusion site-directed mutagenesis kit (Thermo-fisher) yielding in plasmids pJUD36 (encoding C3600-C54S), pJUD37 (encoding C3600-C88S), pJUD38 (encoding C3600-C95S), pJUD39 (encoding C3600-C153S), and pJUD43 (encoding C3600-4C-S). All constructs were verified by DNA sequencing (Eurofins).

### Expression and purification of His_6_-tagged C3600 and variants

His_6_-tagged C3600 variant proteins were expressed in *E. coli* BL21(DE3). Strains were grown in 3L LB supplemented with 100 μg/mL kanamycin until exponential phase, followed by induction with 250 μM isopropyl-β-D-thiogalactopyranoside (IPTG) for 4 hr at 30 ⁰C and 200 rpm. Cells were centrifuged at 8,000 rpm for 10 min, resuspended in 50 mM NaH_2_PO_4_, 300 mM NaCl, 10 mM imidazole (pH 8). After 15 min incubation with 1 mg/ml lysozyme (Goldbio) and 20 ug/ml DNase (Millipore-Sigma), cells were disrupted using a cell disrupter (Constant Systems LTD) and spun down for 30 min at 18,000 rpm at 4 ⁰C. The His_6_-tagged protein variants present in the supernatant were purified using Ni^2+^-NTA affinity chromatography (Goldbio) and dialyzed overnight at 4 ⁰C against 50 mM Tris-HCl (pH 7.5), 200 mM KCl, 0.1 mM MgCl_2_, 0.1 mM EDTA, 1 mM DTT and 10% glycerol (wild-type C3600) or 50 mM potassium phosphate (pH 8), 400 mM NaCl, 2 mM DTT, 1 mM EDTA, and 5% glycerol (C3600-C54S, -C88S, -C95S, and C153S). Proteins were flash-frozen in liquid nitrogen and stored at −80 ⁰C.

### Electrophoretic Mobility Shift Assay (EMSA)

The DNA fragment containing the promoter region of *the c3599-c3600-c3601* operon (P*c3599*) was amplified by PCR using the primer pair listed in Table S4. Purified C3600 wildtype and variant proteins were exchanged into DTT-free buffer (50 mM Tris-HCl buffer [pH 7.5], 200 mM KCl, 0.1 mM MgCl_2_, 0.1 mM EDTA and 10% glycerol) with P-30 gel chromatography columns (Bio-Rad). Increasing concentrations (0.1-1.25 μM) of the C3600 variant proteins were incubated with 2 ng purified P*c3599* in EMSA binding buffer (10 mM Tris-HCl [pH 7.8], 150 mM NaCl, 3 mM magnesium acetate, 10% glycerol, 100 μg/mL bovine serum albumin) for 30 min at room temperature. To study the effect of oxidants on C3600’s DNA binding activity *in vitro*, protein variants were treated with 5-fold molar excess of *N*-chlorotaurine (NCT) or H_2_O_2_ for 10 min. Excess NCT was quenched with 35 μM sodium thiosulfate prior to incubation with P*c3599* DNA fragment. DNA-binding reactions were separated by 6% TBE-polyacrylamide gel electrophoresis, stained with SYBR green (Fisher Scientific) for 30 min in the dark and fluorescence was visualized by UVP ChemStudio Plus (AnalytikJena) and quantified using ImageJ.JS.

### Non-reducing SDS-PAGE

Purified C3600 wildtype and variant proteins were exchanged into DTT-free buffer (50 mM Tris-HCl buffer [pH 7.5], 200 mM KCl, 0.1 mM MgCl_2_, 0.1 mM EDTA and 10% glycerol) with P-30 gel chromatography columns (Bio-Rad). 10 μM proteins were oxidized for 15 min with 5- or 10-molar ratios of HOCl or H_2_O_2_ followed by addition of 1× non-reducing SDS sample buffer. To test reversibility of disulfide bond formation, HOCl-oxidized proteins were treated with 2 mM dithiothreitol (DTT). Proteins were separated by 12% SDS-PAGE and visualized after Coomassie staining.

### Western blot

Overnight MOPSg cultures of BL21(DE3) containing plasmids expressing His_6_-tagged C3600 wild-type, His_6_-tagged C3600-4C-S, or the empty vector control were diluted into fresh MOPSg and incubated at 37 °C and 300 rpm until mid-log phase. When the cultures reached at *A*_600nm_ ∼0.35, protein expression was induced by adding 100 μM IPTG. When cells reached *A*_600nm_ ∼0.55, cultures were either left untreated or treated with 2.5 mM HOCl for 15 min. Cells equivalent to 8 ml of *A*_600nm_= 1 were harvested by centrifugation and incubated in 75 μl of lysis buffer (10 mM KPi [pH 6.5], 1 mM EDTA, 20% [w/v] sucrose, 2 mg/ml lysozyme, 50 U/ml benzonase) supplemented with 0.8 M iodoactamide for 30 min. After a quick freeze thaw cycle and the addition of 360 μl buffer A (10 mM KPi [pH 6.5], 1 mM EDTA), cells disrupted using 0.5 mm glass beads (BioSpec Products) for 30 min at 8 ⁰C. The cell lysates were collected, proteins separated by 12%non-reducing SDS-PAGE and transferred onto a polyvinylidene difluoride (PVDF) membrane (Thermo Fisher Scientific). The membrane was blocked with TBST buffer containing 3% milk powder and 1% BSA, incubated overnight with anti-His antibody (Cell Biolabs) and finally with horseradish peroxidase-conjugated goat anti-mouse IgG (Jackson Immuno-research Laboratory) for 1 h.

## ACKNOWLEDGEMENTS

We thank the Hadjifrangiskou lab (Vanderbilt University) for inviting the Dahl lab to joint lab meetings and the resulting scientific discussions. Dr. Harry Mobley (University of Michigan) and Dr. Shantanu Bhatt (Saint Joseph’s University) are acknowledged for providing strains CFT073 and O127:H6, respectively. This work was supported by Illinois State University School of Biological Sciences startup funds, Illinois State University New Faculty Initiative Grant, and the NIAID grant R15AI164585 (to J.-U. D.). S.S. was supported by Weigel grant by the Phi-Sigma Biological Sciences Honors Society. G.M.A. was supported by the Illinois State University Undergraduate Research Support Program. K. P. H. was supported by a RISE fellowship provided by the German Academic Exchange Service (DAAD). Figures 8 was generated with Biorender.

